# Anti-HIV-1 B cell responses are dependent on B cell precursor frequency and antigen binding affinity

**DOI:** 10.1101/275438

**Authors:** Pia Dosenovic, Ervin E. Kara, Anna-Klara Pettersson, Andrew McGuire, Matthew Gray, Harald Hartweger, Eddy S. Thientosapol, Leonidas Stamatatos, Michel C. Nussenzweig

**Affiliations:** Laboratory of Molecular Immunology, Rockefeller University, 1230 York Avenue, New York, NY 10065-6322; Vaccine and Infectious Disease Division, Fred Hutchinson Cancer Research Center, 1100 Fairview Ave. N., Seattle, WA 98109, USA; Howard Hughes Medical Institute

**Author notes:** Corresponding author: Michel C. Nussenzweig, 1230 York Avenue, New York, NY 10065-6322. Classification: Biological sciences, Immunology and Inflammation.

**Keywords:** B cell, germinal centers, clonal expansion, precursor frequency, antigen affinity, HIV-1, Env

## Abstract

The discovery that humans can produce potent broadly neutralizing antibodies (bNAbs) to several different epitopes on the HIV-1 spike has reinvigorated efforts to develop an antibody based HIV-1 vaccine. Antibody cloning from single cells revealed that nearly all bNAbs show unusual features that could help explain why it has not been possible to elicit them by traditional vaccination, and instead that it would require a sequence of different immunogens. This idea is supported by experiments with genetically modified immunoglobulin knock-in mice. Sequential immunization with a series of specifically designed immunogens was required to shepherd the development of bNAbs. However, knock-in mice contain super-physiologic numbers of bNAb precursor expressing B cells and therefore how these results can be translated to a more physiologic setting remains to be determined. Here we make use of adoptive transfer experiments using knock-in B cells that carry a synthetic intermediate in the pathway to anti-HIV-1 bNAb development to examine how the relationship between B cell receptor affinity and precursor frequency affects germinal center B cell recrutiment and clonal expansion. Immunization with soluble HIV-1 antigens can recruit bNAb precursor B cells to the germinal center when there are as few as 10 such cells per mouse. However, at low precursor frequencies the extent of clonal expansion is directly proportional to the affinity of the antigen for the B cell receptor, and recruitment to germinal centers is variable and dependent on re-circulation.

**Significance statement:** An essential requirement for an HIV-vaccine is to elicit antibodies to conserved regions of the spike protein (Env) becasue these antibodies can protect against infection. Although broadly neutralizing antibodies develop naturally in rare individuals after prolongued HIV infection, eliciting them by vaccination has only been possible in artificial knock-in mouse models wherein the number of B cells expressing the antibody precursor is super-physiologic. To understand the relationship between precursor frequency, antigen affinity and germinal center recruitment we have performed adoptive transfer experiments in which fixed numbers of precursor cells are engrafted in wild type mice. Our results provide a framework for understanding how precursor frequency and antigen affinity shape humoral immunity to HIV.

## Introduction

Most vaccine responses are elicited by immunization with either attenuated, heat killed or genetically inactivated pathogens, or pathogen-derived proteins (1, 2). High affinity antibodies are the key effectors of protection in the great majority of these vaccines (3, 4). These antibodies develop in germinal centers, which are the microanatomic site of B cell clonal expansion and antibody affinity maturation (5).

Although there is no effective vaccine against HIV-1, potent broadly neutralizing antibodies (bNAbs) that bind to HIV-1 envelope glycoprotein (Env) with high affinities can prevent infection in animal models even when present at low concentrations (6-9). However, elicitation of bNAbs to HIV-1 Env presents a series of what may be unique problems including a requirement for one or more of the following unusual features: high levels of somatic hypermutation, incorporation of self-glycans into a complex peptide-glycan epitope, long CDRH3s, and self-or poly-reactivity (10-13). Among these the requirement for higher than normal levels of somatic mutation, a reaction that occurs in germinal centers, is nearly universal (14).

B cell entry into and persistence in germinal centers is dependent on a large number of variables including but not limited to: the presence or absence of T cell help, antigen concentration, B cell receptor affinity for the antigen, responding B cell precursor frequency, the valency of the antigen, and the physiological state of the B cell (5, 15, 16). The role of B cell receptor affinity and precursor frequency in B cell entry into germinal center responses has been studied in wild type (WT) mice (17-20). For example, germinal centers in WT mice immunized with the hapten (4-hydroxy-3-nitrophenyl)-acetyl (NP) are colonized by B cells with highly varied affinities, some as low as 500 μM (*K*_*a*_) (18). Similar results have also been found in WT mice immunized with *Bacillus* anthracis protective antigen, influenza hemagglutinin and chicken gamma globulin (19, 20).

In contrast to WT mice, results using Ig transgenic or knock-in B cells, with defined affinities for their cognate antigen, display far more variable results. Transgenic or knock-in B cells that carry anti-NP antibodies with affinities as low as 300 μM enter germinal centers and undergo affinity maturation unless outcompeted by higher affinity antibody expressing B cells (21-24). However, knock-in B cells that carry an anti-hen egg lysozyme (HEL)-specific receptor require high affinity interactions (< 23 μM) for efficient expansion (25, 26). Similarly, B cells that carry anti-HIV-1 Ig knock-in genes also appear to require high affinity interactions with polymerized antigen to gain access to and participate in germinal center reactions (27-30).

Whether the high affinity requirements found in the anti-HIV-1 Ig knock-in mice represent a general rule for HIV-1-specific B cells, or alternatively whether lower affinity interactions characteristic of B cells participating in germinal center reactions under physiologic conditions can induce HIV-1-specific B cell responses has not been determined. This question is particularly important for vaccines aimed at eliciting bNAbs to HIV-1 because they appear to require the recruitment of rare B cell precursors into germinal centers and sequential exposure to different immunogens (31-35).

Here we examine the relationship between precursor frequency and antigen binding affinity for B cells that carry a knock-in B cell receptor specific for the HIV-1 CD4-binding site.

## Results

### B cell development in 3BNC60^SI^ knock-in mice

To study the relationship between B cell precursor frequency, affinity, and epitope-specific B cell responses to HIV-1 Env we sought to use B cells that develop normally, display nearly complete allelic exclusion and show normal levels of cell surface IgM and IgD. This is particularly important for anti-HIV-1 IgH + IgL knock-in mice because a number of these mice, including 2F5 (29, 36, 37), 4E10 (38), 3BNC60 germline (30), and VRC01germline (28), display combinations of abnormal B cell development, anergy, and/or absence of allelic exclusion.

B cell development in the bone marrow of 3BNC60^SI^ knock-in mice, expressing a synthetic intermediate antibody composed of the mature 3BNC60 IgH, and the germline IgL (30, 31), showed the expected decreases in pre- and immature B cells, and increases in mature follicular B cells that are associated with expression of a non-autoreactive pre-rearranged knock-in B cell receptor (Figure 1A and B) (15, 39-41). Similarly, the spleen contained decreased levels of transitional T1 and T3 B cells, characteristics found in non self-reactive Ig transgenic mice while mice that carry self-reactive Igs show increased frequencies of T3 B cells (42-44). 3BNC60^SI^ knock-in mice also displayed increased frequencies of splenic follicular and marginal zone B cells (Figure 1C and D). Thus, B cells in the bone marrow and spleen of 3BNC60^SI^ knock-in mice display no signs of self-reactivity or anergy, as measured by the size of B cell subpopulations, and IgM/IgD expression levels respectively.

**Figure 1.**
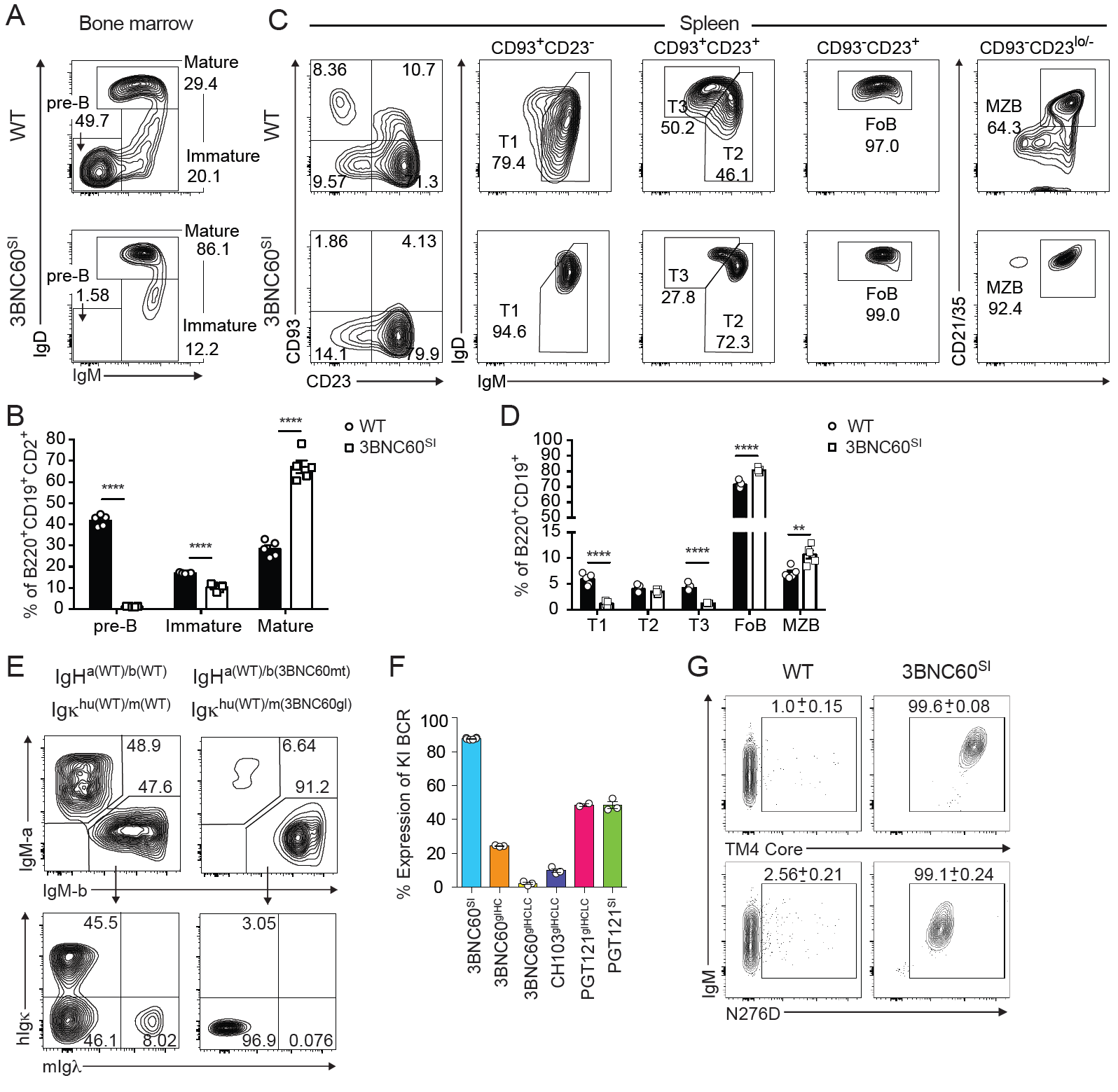
B cell development in 3BNC60^SI^ knock-in mice. **(A and B)** Representative flow cytometric analysis (**A**) and quantitation (**B**) of B cell development in bone marrow of wild type C57Bl/6 (WT) and 3BNC60^SI^ knock-in mice. Pre-B cells (IgM^-^IgD^-^), immature- (IgM^+^IgD^-^) and mature B cells (IgM^+/lo^IgD^+^) are shown (n=5 mice per genotype; mean±SEM). Plots are pre-gated on B220^+^CD19^+^CD2^+^ cells. **(C and D)** Representative flow cytometric analysis (**C**) and quatitation (**D**) of B cell populations in spleen of WT and 3BNC60^SI^ knock-in mice. Transitional (T) 1 (CD93^+^CD23^-^IgM^+^), T2 (CD93^+^CD23^+^IgM^hi^IgD^+^), T3 (CD93^+^CD23^+^IgM^lo^IgD^hi^), follicular (FoB; CD93^-^ CD23^+^IgM^+^IgD^hi^) and marginal zone (MZB; CD93^-^CD23^lo^IgM^+^CD21/35^+^) B cells are shown (n=5 mice per genotype; mean±SEM). Plots are pre-gated on B220^+^CD19^+^ cells. **(E)** Frequency of IgH-a^+^ and IgH-b^+^ B220^+^ cells (upper panels) and the frequency of human (h) Igκ constant and mouse (m) IgΛ amongst IgH-b^+^B220^+^ cells (lower panels) in heterozygous WT (left panels) and 3BNC60^SI^ knock-in (right panels) mice crossed to IgH-a, hIgκ mice (mt = mature, gl = germline). Representative flow cytrometic graphs of WT (n=3) and 3BNC60^SI^ knock-in (n=6) mice are shown **(F)** Frequencies of B cells that express the knock-in (KI) B cell receptor (BCR; IgH-b^+^hIg^κ-^mIgΛ^-^) from the heterozygous offspring of anti-HIV-1 Ig knock-in mice (indicated on *x*-axis) crossed to IgH-a, hIgκ mice (n=2-6 mice per genotype; mean±SEM; HC = heavy chain, LC = light chain). **(G)** Representative flow cytometric analysis of TM4 core (high affinity antigen) or N276D (intermediate affinity antigen) binding B220^+^IgM^+^ cells from WT (n=3) and 3BNC60^SI^ knock-in (n=11) mice. Numbers in plots indicate the mean±SEM frequency of antigen-binding B cells. ^**^, P < 0.01; ^****^, P < 0.0001. Two-tailed unpaired Student’s *t*-test.

Absence of allelic exclusion is associated with self-reactivity or otherwise abnormal B cell receptors (45). To evaluate allelic exclusion and receptor editing, we combined IgH^a/b^ allotypes with a heterozygous human Ig kappa constant region knock-in (hIgκ) allele (46). Due to allelic exclusion, ∼50% of WT B cells express either IgH^a^ or IgH^b^. Of the IgH^b^ or IgH^a^ expressing cells, ∼45% express hIgκ and ∼45% mouse (m)Igκ, while the remaining 5-10% express mIgΛ light chains (Figure 1E, left panels). In contrast, and as expected from non self-reactive pre-recombined knock-in Igs, 90% of heterozygous 3BNC60^SI^ knock-in B cells express the knock-in IgH^b^ allele in combination with the 3BNC60 Igκ light chain (Figure 1E, right panels). This is in contrast to numerous other strains of anti-HIV-1 Ig knock-in mice that we and others have generated that show far lower levels of knock-in BCR expression (Figure 1F and (28, 36, 38)). In addition, 99.6% and 99.1% of circulating B cells from homozygous 3BNC60^SI^ knock-in mice bound high (TM4 core) and intermediate (N276D) affinity HIV-1 Env proteins respectively, as compared to 1.0 and 2.6% in WT controls. Furthermore, the magnitude of IgM expression by Env-binding 3BNC60^SI^ knock-in B cells is comparable to WT indicating that these cells are not anergic (39) (Figure 1G). In summary, 3BNC60^SI^ knock-in B cells show efficient allelic exclusion of WT alleles and are therefore relatively homogenous in terms of their antigen binding properties. Moreover, 3BNC60^SI^ knock-in B cells show no appreciable correlates of self-reactivity.

### Relationship between precursor frequency and clonal expansion

To track the fate of antigen stimulated epitope-specific precursor B cells, allotype marked 3BNC60^SI^ knock-in B cells (CD45.2) were transferred into congenic (CD45.1) WT mice. As reported by others (17), the efficiency of transfer was approximately 5% as determiend by dilution experiments (Supplementary Table 1). To determine whether 3BNC60^SI^ knock-in B cells are responsive to soluble HIV-1 Env, we immunized WT mice engrafted with 100,000 3BNC60^SI^ knock-in B cells with soluble N276D Env which binds to the knock-in BCR with ≈ 40 μM affinity (Figure 2A and Supplementary Table 2). Three days after immunization, an average of 0.3% of all B cells in both naive and immunized recipient mice were knock-in cells (Figure 2B). Germinal centers in draining lymph nodes peaked as early as 7 days after immunization (Figure 2C), when the knock-in cells had expanded to represent 4-10% of all B cells (Figure 2B, closed circles) and over 60% on average of all B cells in GCs (Figure 2D), compared to non-immunized recipients where there was no such expansion (Figure 2B, open circles). We conclude that when 100,000 3BNC60^SI^ knock-in B cells are engrafted into WT mice they can develop into GC B cells and proliferate extensively after stimulation with soluble antigen that binds to the B cell receptor with ≈ 40 μM affinity (Figure 2B-E).

**Figure 2.**
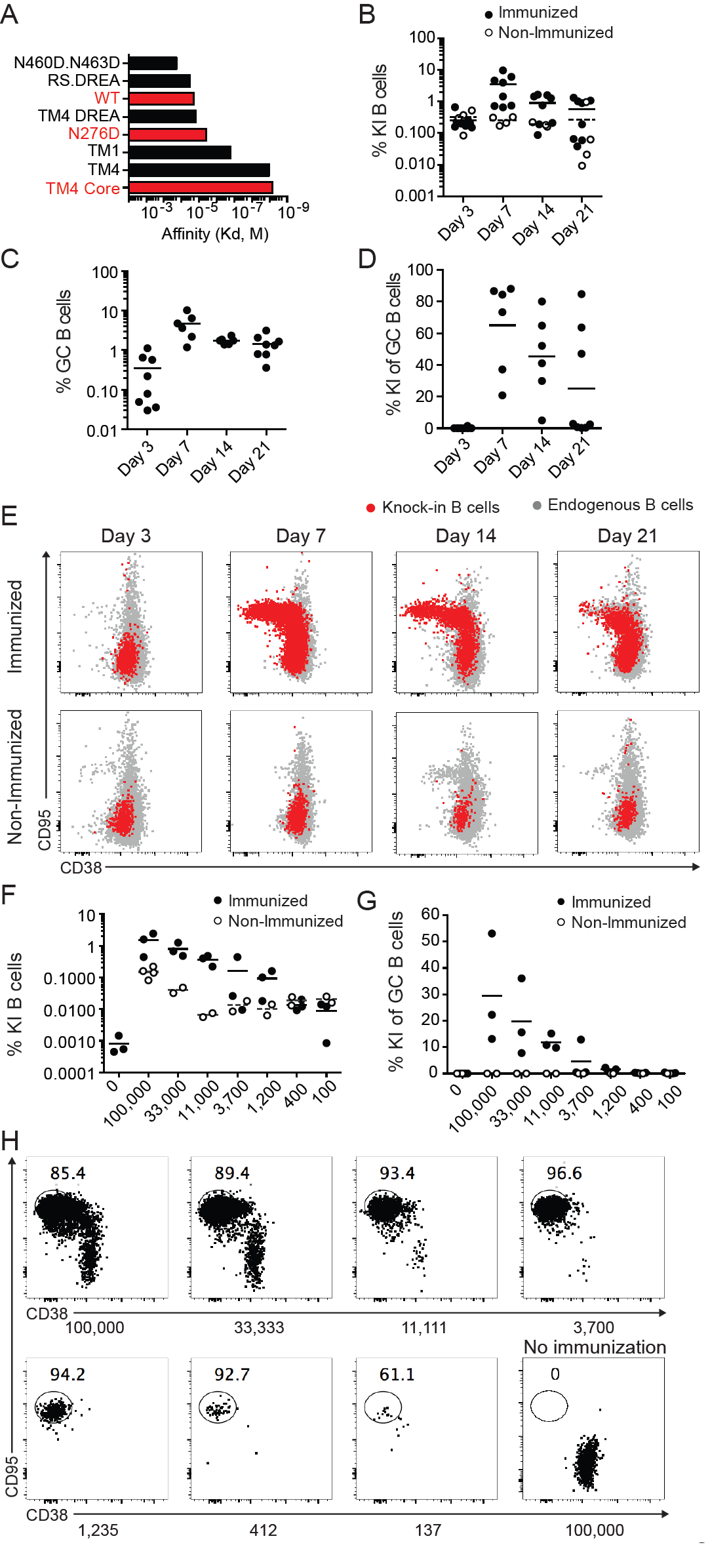
3BNC60^SI^ knock-in B cells transferred into WT recipient mice expand and form germinal centers following soluble HIV-I Env immunization. **(A)** Bio-Layer Interferometry (BLI) analysis of the affinity for selected 426c-Env-based trimers against 3BNC60^SI^ Fab’. Numbers on the *x*-axis refer to individual proteins listed in the legend. **(B-E)** 10^5^ 3BNC60^SI^ knock-in (KI) B cells (CD45.2) were engrafted into recipient B6.SJL (CD45.1) mice that were subsequently immunized with soluble N276D Env trimers (see also Supplementary Table 2). (**B**) Frequency of 3BNC60^SI^ knock-in B cells in immunized (closed circles) and non-immunized (open circles) recipient mice. (**C**) Total GC B cells of immunized recipient mice (**D**) Frequency of 3BNC60^SI^ knock-in B cells amongst GC B cells of immunized recipient mice. (**E**) Representative flow cytometric analysis of GC B cells (Fas^+^CD38^-^) amongst transferred 3BNC60^SI^ knock-in B cells (red) and endogenous B cells. Data in B-E are pooled from two independent experiments (n=2-8 mice per time point). Frequency of 3BNC60^SI^ knock-in B cells (**F**) and per GC B cells (**G**) in recipient mice engrafted with the indicated numbers of 3BNC60^SI^ knock-in B cells (below) 14 days following soluble N276D Env immunization (closed circle). Mice receiving the same number of cells with no immunization are shown for comparison (open circles). **(H)** Representative flow cytometric analysis of mice in **F and G**. Percentage of GC B cells amongst 3BNC60^SI^ knock-in B cells are shown in plots (see also Supplementary Figure 1). Data in F-H are representative of two independent experiments (n=2-3 mice per group per time point). Bars in all graphs indicate mean values.

To examine the relationship between precursor frequency and B cell expansion in response to a soluble HIV-1 Env antigen we engrafted varying numbers of 3BNC60^SI^ knock-in B cells into WT recipients and analyzed the B cell responses 14 days after immunization. Mice engrafted with 1,200-100,000 B cells showed 10-50 fold increases in the fraction of 3BNC60^SI^ knock-in B cells after immunization with soluble N276D Env (≈ 40 μM) (Figure 2F). In mice engrafted with a high number of KI cells, these cells contributed up to 50% of all B cells in GCs (Figure 2G). Moreover, engraftment with as few as 100 3BNC60^SI^ knock-in B cells was sufficient for their recuritment into the GC after immunization with a soluble ≈ 40 μM affinity antigen (Figure 2H and Supplementary Figure 1). We conclude that immunization with a soluble antigen of modest affinity expands epitope-specific precursor B cells in a manner that is directly correlated to the number of epitope-specific precursors present at the time of immunization.

### Relationship between affinity, precursor frequency and clonal expansion

To examine the role of HIV-1 Env antigen affinity in 3BNC60^SI^ knock-in B cell responses to immunization we engrafted WT mice with 10,000, 100 or 10 cells and immunized with either of the three soluble Env trimer antigens of varying affinities (≈ 7nM, ≈ 40 μM and ≈ 200 μM, Figure 2A and Supplementary Table 2). The frequency of knock-in B cells in the draining lymph nodes and their phenotype was determined 14 days after immunization. In the absence of immunization transferred B cells were only detected in mice engrafted with 10,000 cells and they remained naïve (CD95^-^CD38^+^, Figure 3A).

**Figure 3.**
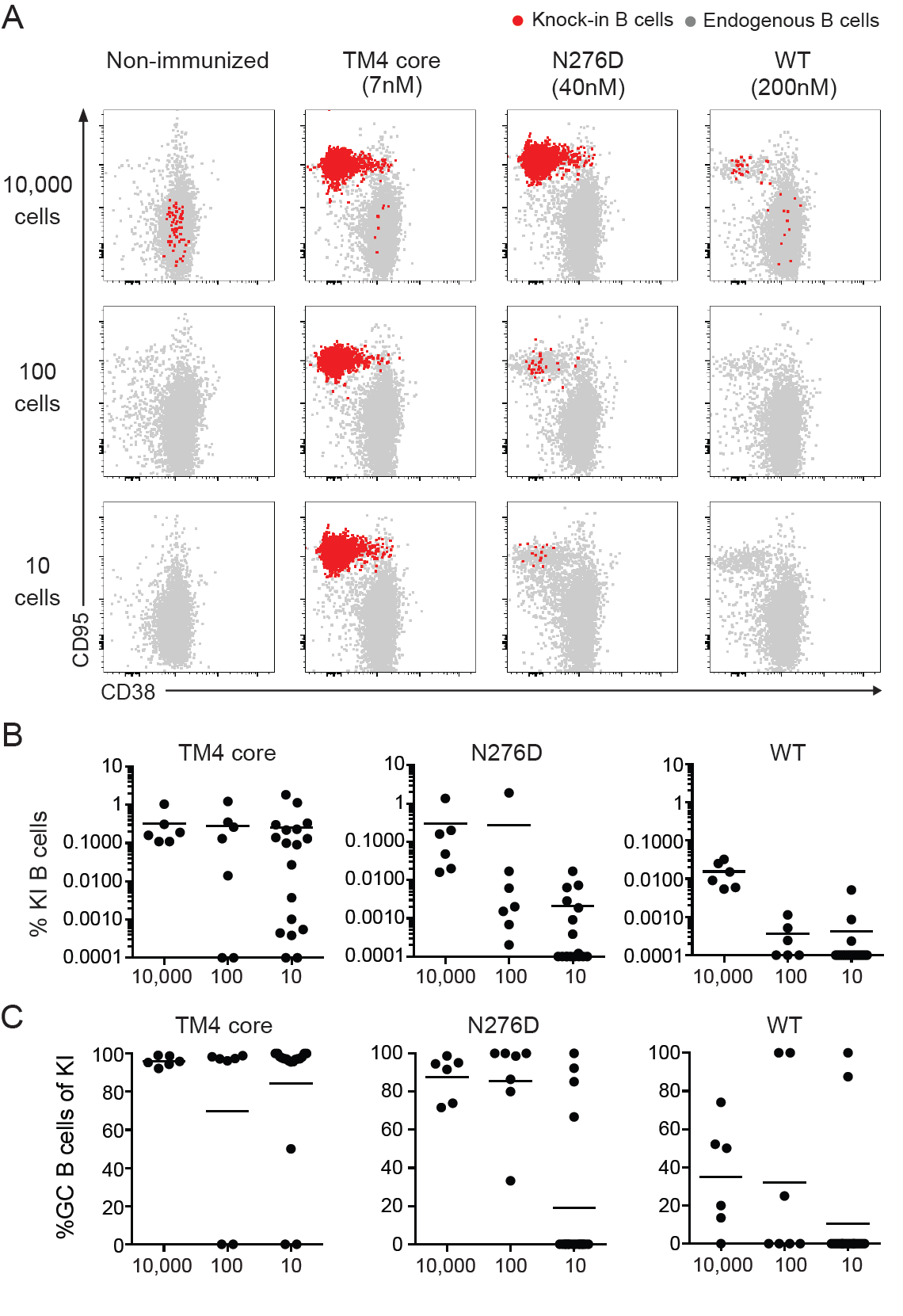
Relationship between antigen affinity and precursor frequency. **(A)** Representative flow cytometric analysis of GC B cells in B6.SJL (CD45.1) recipient mice engrafted with 10 000 (n=6 mice/antigen), 100 (n=8 mice/antigen) or 10 (n=20 mice/antigen) 3BNC60^SI^ knock-in (KI) B cells (CD45.2) 14 days following immunization with the indicated soluble Env trimer protein (top, see also Supplementary Table 2). 3BNC60^SI^ knock-in (red) and endogneous (gray) GC B cell responses are shown. (**B**) Frequency of 3BNC60^SI^ knock-in B cells from mice in A. Animals showing undetectable frequencies of knock-in B cells were set to 0.001 to indicate the number of non-responders. (**C**) Frequency of GC B cells amongst 3BNC60^SI^ knock-in B cells from mice in A. Data are pooled from two independent experiments. Bars in all graphs indicate mean values.

All antigens produced similar sized GCs in the control mice that did not receive 3BNC60^SI^ knock-in B cells indicating similar overall immunogenicity by the different antigens (Supplementary Figure 2). Mice immunized with high affinity soluble antigen (≈ 7nM) showed high levels of 3BNC60^SI^ knock-in B cell expansion irrespective of the number of precursors present at the time of immunization (Figure 3A and B). In addition, nearly all of the transferred B cells in the draining lymph nodes of these mice diplayed a GC B cell phenotype (CD95^+^CD38^-^) (Figure 3C). However, the number of mice with precursor cells that responded and the frequency of B cell expansion was variable in mice engrafted with smaller numbers of precursors. As described above (Figure 2B-H), 3BNC60^SI^ knock-in B cells also responded to immunization with the intermediate affinity soluble antigen (≈ 40 μM) even when present at levels as low as 10 precursor B cells per mouse. However, the degree of B cell expansion and their relative contribution to the GC reaction was decreased and variable in comparison with the high affinity antigen. Finally, even immunization with the low affinity soluble antigen (≈ 200 μM) produced GC responses by the engrafted cells, but this response was highly variable and mainly found in mice engrafted with the highest number of precursor cells (Figure 3A-C).

The difference in the relative amount of B cell expansion in response to antigens of differing affinities could be seen as early as 3-4 days after immunization. Engrafted 3BNC60^SI^ knock-in B cells labeled with Celltrace violet (CTV) diluted the dye to a greater extent in mice immunized with high affinity than with intermediate affinity antigen, where no cell division was detected at this time point (Supplementary Figure 3 A and B). Thus, in the context of a polyclonal immune system where a given precursor is present in only small numbers, a higher affinity antigen is essential for reproducible induction of high levels of epitope-specific B cell expansion. Nevertheless, even lower affinity soluble antigens can recruit B cell precursors present in limiting numbers to the GC reaction, but this response is more variable than when large number of precursors are present.

### Rare precursor B cell expansion is dependent on lymph node homing

Our analysis is limited to the draining lymph nodes, and the probability that a rare engrafted cell is present in those nodes at the time of immunization is small. Thus, it is possible that under conditions of limiting precursor frequency, GC participation requires that engrafted 3BNC60^SI^ knock-in B cells survey the animal and enter lymph nodes containing ongoing GC reactions even after the initiation of the reaction (24, 47). To examine this possibility we blocked lymphocyte homing and egress from lymph nodes with a combination of a neutralizing antibody to CD62L, and FTY-720 (S1PR1 antagonist). When compared to controls engrafted with 10 3BNC60^SI^ knock-in B cells, anti-CD62L and FTY-720-treated mice immunized with the high affinity (7nM) soluble antigen showed significantly decreased levels of precursor B cell expansion (Figure 4A) and GC B cell entry (Figure 4B and C). We conclude that the efficient activation of rare naïve precursor cells with a high affinity antigen is dependent on the recirculation of naïve B cells.

**Figure 4.**
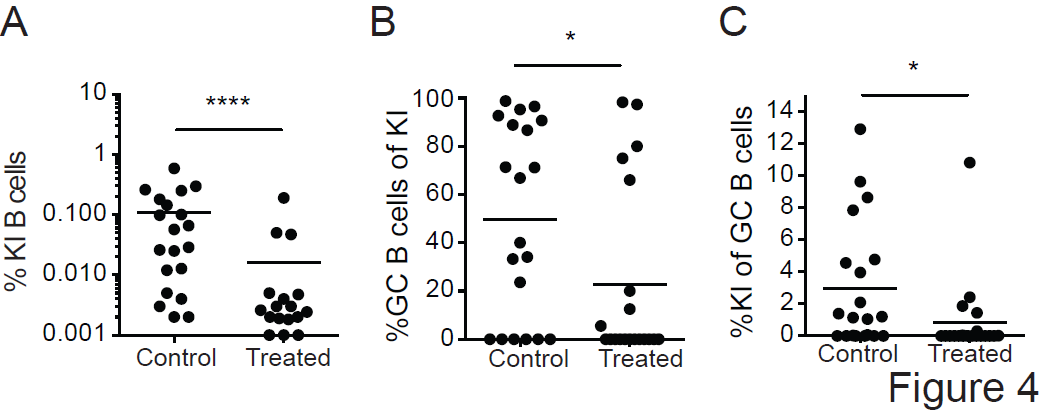
Recruitment of rare precursors to germinal centers is dependent on B cell recirculation. 10 3BNC60^SI^ knock-in (KI) B cells (CD45.2) were engrafted into recipient B6.SJL (CD45.1) mice and immunized with soluble TM4 core (high affinity Env, ≈ 7 nM). To block homing and egress of lymphocytes, recipient mice were treated with neutralizing antibody to CD62L and FTY-720 (treated; n=20 mice) on the day of immunization and every 48 h thereafter until the day of analysis (day 14). Control mice were treated with Rat IgG2a isotype antibody and vehicle (control; n=20 mice). Frequency of 3BNC60^SI^ knock-in B cells (left), frequency of GC B cells amongst 3BNC60^SI^ knock-in B cells (middle panel), and frequency of 3BNC60^SI^ knock-in B cells amongst GC B cells (right panel). Data are pooled from two independent experiments. ^*^, P < 0.05; ^****^, P < 0.0001. Two-tailed unpaired nonparametric Mann-Whitney U-test. Bars in all graphs indicate mean values.

## Discussion

Potent bNAbs to HIV-1 Env are protective in animal models even at low concentrations, and it is generally accepted that a vaccine that elicits such antibodies would be efficacious (6-9, 12, 48). However, these antibodies have only been elicited by immunization in mice that carry human Ig knock-in genes using a sequence of immunogens specifically designed to target the knock-in B cell receptor (33-35). Although these experiments establish that sequential immunization can produce bNAbs from specific germline precursors, there are several important issues that must be resolved before these results can be translated to genetically un-manipulated animals and humans. Among these are the requirements for antigen affinity in recruiting anti-HIV-1 bNAb precursor cells to germinal centers, and how affinity requirements relate to precursor frequency, and the clonal expansion of the target cell.

To date, experiments with knock-in mice expressing the 2F5 and germline VRC01 anti-HIV-1 antibodies suggest a requirement for high affinity polymerized antigen for B cell clonal expansion (27-29). 2F5 knock-in mice carry a self-reactive receptor that fails to support B cell development and only small numbers of cells escape central tolerance and enter the periphery with an anergic phenotype (27, 29, 36, 37). Nevertheless, these self-reactive B cells can expand when the knock-in mice are immunized with a high affinity polymerized antigen (27, 29). Similar results have also been obtained with the model antigen HEL using B cells from mice that express HEL as a neo-self antigen and carry a self-reactive anti-HEL antibody gene (49).

A requirement for high affinity (0.5μM) polymerized antigens was also found in adoptive transfer experiments using germline VRC01 knock-in B cells (28). Moreover, high levels of B cell expansion were only found when specific precursors were present at concentrations of 1:10^5^ or 200-400 cells per mouse. One potential explanation for the difference between VRC01 knock-in B cells and the results obtained in studies of germinal centers in WT mice (18-20) is that VRC01 knock-in B cells display some features of abnormal B cell development (28). Heterozygous VRC01 knock-in B cells fail to show high levels of allelic exclusion. They co-express knock-in Igκ and endogenous IgΛ light chains (28). Under physiologic circumstances only 2-5% of B cells express dual light chain receptors (46, 50). Ig transgenic or knock-in mice like 3BNC60^SI^ that express non-self-reactive receptors also show the same high levels of allelic exclusion (46, 50-52). In contrast, dual expressing B cells like VRC01 knock-in where an unusually large proportion of B cells displayed double expression of κ and Λ light chains (28) are typically found in autoimmune B cells of mice and humans (45, 53-56). These cells arise by receptor editing, whereby B cells expressing self-reactive receptors, or incompatible Ig heavy and light chains have their receptors replaced or diluted by continuing V(D)J recombination (39, 46, 57-60). Transgenic or knock-in B cell receptors that are designed to be resistant to this process can nevertheless be edited by co-expressing a second light chain, as seen in VRC01 knock-in B cells (28). Co-expression of a second light chain produces chimeric receptors, which dilute the effects of the receptor that fails to support normal B cell development (39, 46, 57-60). In such cases each developing B cell expresses a unique and random combination of receptors composed of the knock-in Ig heavy and either/or the knock in light chain and endogenous light chains. The result is that many of the developing B cells fail to bind to their cognate antigen and those that do so may display variable levels of affinity and avidity.

Our experiments were performed using 3BNC60^SI^ knock-in B cells that show normal levels of allelic exclusion, and the expected uniform levels of antigen binding. In addition, 3BNC60^SI^ knock-in B cells show no measurable signs of anergy or self-reactivity. The data indicate that even when present in limiting numbers (10 precursor B cells per mouse) such cells can be recruited to germinal centers by soluble HIV-1 Env antigens. Soluble high affinity antigen (≈ 7 nM) induced high levels of clonal expansion in draining lymph nodes when as few as 10 specific precursor cells were present in the mouse. Although lower affinity antigens (≈ 40-200 μM) were able to recruit 3BNC60^SI^ knock-in B cells into the germinal center, B cell expansion was limited and their recruitment was more variable. Our results are in keeping with observations made in anti-NP knock-in and WT mice in that B cells found in germinal centers of mice immunized with the hapten NP, or with protein antigens can display micromolar affinities for the immunogen (18-20). The differing levels of expansion induced by antigens of varying affinity are likely related to B cell competition for T cell help at the T-B border and within germinal centers (19, 23, 61-63).

3BNC60^SI^ differs from 2F5 and VRC01 in that it is not a germline precursor but a synthetic intermediate in HIV-1 bNAb development composed of a mutated heavy chain and a germline light chain. Moreover, B cells in 3BNC60^SI^ appear to develop normally, whereas several different lines of anti-HIV-1 Ig knock-in mice show abnormalities associated with self-or polyreactivity including germline 2F5, VRC01, 3BNC60, CH103, and to a lesser extent PGT121 (Figure 1F and (28, 30, 33, 36, 37)). The abnormalities in development found in these knock-in mice are entirely consitent with the observation that anti-HIV-1 antibodies are frequently self- and/or poly-reactive (10, 64, 65). Whether or not B cells expressing authentic bNAb precursors in humans show the same developmetal defects and differ from B cells that develop normally in their requirements for germinal center recruitment and clonal expansion remains to be determined.

## Materials and methods

### Mice

3BNC60^SI^ (<u>s</u>ynthetic intermediate) knock-in mice carry the pre-rearranged *Ig* V(D)J genes encoding the mature heavy chain (somatically mutated) and predicted germline light chain of human bNAb 3BNC60 (30, 31). Mice used in experiments were homozygous for both knock-in alleles (CD45.2). IgH^a/a^Igκ ^hu/hu^-, PGT121 germline (glHCLC) and PGT121^SI^ were described previously (33, 46). CH103 germline (glHCLC) knock-in mice were generated as described previously (31, 33) C57Bl/6 wild type (WT) and B6.SJL (CD45.1) mice were obtained from Jackson Laboratories. All experiments were conducted with approval from the Institutional Review Board and the IACUC at The Rockefeller University.

### HIV-1 Env antigens

Soluble trimeric gp140 Envs were expressed in 293 F cells and purified as previously described (30). All are derived from the Clade C 426c gp140 Env (66) or 426c TM4ΔV1-3 (herein called TM4 Core) (31). Briefly, expression plasmids encoding Env proteins were transfected into 293 F cells at a density of 10^6^ cells ml^-1^ in Freestyle 293 media (Life Technologies) using the 293Free transfection reagent (EMD Millipore). Expression was carried out in Freestyle 293 media for 6 days with gentle shaking at 37 °C in the presence of 5% CO_2_ after which cells and cellular debris were removed by centrifugation at 10,000*g* followed by filtration through a 0.2 μM filter. Clarified cell supernatant was passed over Agarose-bound Galanthus Nivalis Lectin (GNL) resin (Vector Laboratories), pre-equilibrated with 20 mM Tris, 100 mM NaCl, 1 mM EDTA pH 7.4 (GNL binding buffer), followed by extensive washing with GNL binding buffer. Bound protein was eluted with GNL binding buffer containing 1 M methyl mannopyranoside. The eluted protein was run over 16/60 S200 size-exclusion column (SEC) pre-equilibrated in PBS. Fractions containing trimeric gp140 protein were pooled, aliquoted, frozen in liquid nitrogen and stored at -80 °C.

Monomeric TM4 core (amino acids 44-494 HXB2 numbering) with a C terminal hexa-His and Avi-Tag was expressed, purified, and biotinylated as previously described (30, 31). Avitagged 426c.NLGS.TM4.ΔV1-3 was biotinylated *in vitro* using the *In Vitro* Biotin Ligase Kit (Avidity), followed by SEC using a 10/300S 200 column (GE Healthcare) equilibrated in PBS to remove unligated biotin and BirA enzyme.

### BLI assay

BLI assays were performed on the Octet Red instrument (ForteBio, Inc, Menlo Park, CA) as previously described (30). gp140s were biotinylated using EZ-Link NHS-PEG4-Biotin (Thermo Scientific) at a ratio of one biotin molecule per gp140 trimer (0.33 biotin molecules per monomer). Unligated biotin was removed using Zebra desalting columns (Thermo Scientific). Biotinylated trimeric recombinant gp140s were immobilized on streptavidin biosensors (ForteBio) at concentrations that yielded the same *R*_max_ for all Envs tested (1–2 μM). The baseline signal was recorded for 1 min in KB, then the sensors were immersed into wells containing dilutions of purified recombinant Fabs (4–0.125 μM) for 5 min (association phase). The sensors were then immersed in KB without Env for an additional 10 min (dissociation phase). Curve fitting was performed using a 1:1 binding model and the Data analysis software (ForteBio). Mean *k*_on_ and *k*_off_ and *K*_A_ values were determined by averaging all binding curves that matched the theoretical fit with an *R*^2^ value of ⩾ 0.95. All measurements of Env-Ab binding were performed at 30 °C with shaking at 1,000 r.p.m and corrected by subtracting the signal obtained from simultaneous traces performed with the corresponding envelope proteins in the absence of antibody, using PBS only.

### Lymphocyte preparation

Lymphocytes were harvested by forcing lymph nodes and/or spleen through 70 μm filters (BD) into RPMI 1640 media (Gibco) containing 6 % fetal bovine serum (FBS) and 10 mM HEPES, followed by lysis of erythrocytes using 1X ACK Lysis Buffer (Gibco). B cells were enriched by negative selection using anti-CD43 MicroBeads (Militenyi Biotec) with magnetized LS columns according to manufacturers’ instructions. In some experiments, enriched B cells were labelled with Celltrace violet (CTV; Thermo Scientific) according to manufacturers’ instructions.

### Animal experiments

Enriched B cells were transferred intravenously into B6.SJL recipient mice. 10 μg of indicated soluble Env proteins were injected subcutaneous in Imject Alumn (Thermo Scientific). To block homing and egress of lymphocytes into lymph nodes, mice were administered neutralizing antibody to CD62L (clone Mel-14; 100 μg per dose; BioXCell) and FTY-720 (3 mg/kg per dose; Cayman Chemical) intraperitoneally at the time of immunization and every other day until the day of analysis. Control mice were treated with Rat IgG2a (clone 2A3; 100 μg per dose; BioXCell) and vehicle (physiological saline 0.9% NaCl; Sigma).

### Flow cytometry

Single cell suspensions of lymphocytes were maintained at 4 ºC in FACS buffer (PBS containing 2 % FBS and 1 mM EDTA). Fc receptors were blocked using rat anti-mouse CD16/32 (clone 2.4G2; BD) before staining with fluorophore-conjugated antibodies to: B220, CD19, CD23, CD21/35, CD93, CD2, IgM, IgD, CD38, CD45.1, CD45.2 (eBioscience), CD95, CD4, CD8, NK1.1, Gr1 or F4/80 (BD). Dump staining including CD4, CD8, NK1.1, Gr1 and F4/80 were included in all flow cytometry stainings. Dead cells were excluded from analyses using Zombie NIR^(tm)^ Fixable Viability Kit (BioLegend) or LIVE/DEAD^(tm)^ Fixable Aqua Dead Cell Stain (Invitrogen) reagents. Env-specific B cells were stained using Avi-tagged DMRScore-or N276D-gp120 monomers (5 μg/ml) and detected using Streptavidin-PE (BD). Samples were aquired on a BD Fortessa and analyzed using FlowJo software (Treestar).

### Determination of B cell transfer efficiency

A defined number of knock-in B cells (CD45.2) were transferred into B6.SJL recipient mice (CD45.1). The next day, spleen and peripheral lymph nodes were collected and absolute number of lymphocytes were calculated using a hemocytometer. Samples were next analyzed by flow cytometry to determine the frequency of knock-in B cells present in the sample. This information was used to determine the absolute number of knock-in B cells present in recipient mice at the time of immunization.

### Statistics

Data were analyzed with Prism 6 (GraphPad) using two-tailed unpaired Student’s *t*-tests (normally distributed data sets comparing the mean of two samples) or two-tailed nonparameteric Mann-Whitney U-tests (data sets that were determined by an *F* test not to have a normal distribution and where there was a comparison of the mean of two samples) as indicated in-text. For all analyses, *P* ≤ 0.05 was considered statistically significant.

## Acknowledgements

We thank Thomas Eisenreich and Steven Tittley for animal husbandry; Zoran Jankovic for laboratory support; and Mila Jankovic for comments on the manuscript. This work was supported in part by grants: R01AI081625 (L.S), R01 AI104384(L.S), the Bill and Melinda Gates Foundation Collaboration for AIDS Vaccine Discovery (CAVD) grants OPP1092074 and OPP1124068 (M.C.N.), NIH Center for HIV/AIDS Vaccine Immunology and Immunogen Discovery (CHAVI-ID) 1UM1 AI100663-06 (M.C.N.), and NIH grant AI037526-24 (M.C.N). P. D. is supported by an NIH award (K99AI127243-01A1); E.E.K. is supported by a National Health and Medical Research Council C.J. Martin Overseas Biomedical Fellowship; H.H. is supported by the Cancer Research Institute Irvington Fellowship. M.C.N. is a Howard Hughes Medical Institute Investigator.

---

**Supplementary Figure 1. Percent of 3BNC60^SI^ knock-in B cells in GCs 14 days after immunization with soluble N276D Env trimers.** Frequency of GC B cells amongst 3BNC60^SI^ knock-in B cells 14 days after immunization with soluble N276D Env. Number of engrafted 3BNC60^SI^ knock-in B cells at the time of immunization is indicated on the *x*-axis. Data are representative of two independent experiments (n=2-3 mice per group per time point)

**Supplementary Figure 2. Percent GC B cells in wild type mice after immunization with selected soluble Env antigens.** Frequency of GC B cells in wild type mice 14 days after immunization with indicated soluble Env proteins. Data are pooled from two independent experiments (n=4-7 mice per time point). Statistics were calculated using ordinary one-way ANOVA (Tukey). ns = not significant (p>0.05)

**Supplementary Figure 3. Proliferation of engrafted 3BNC60^SI^ knock-in B cells 4 days after soluble Env immunization. (A)** Histograms show cell trace violet (CTV) dilution profiles of 10,000 3BNC60^SI^ knock-in B cells engrafted into B6.SJL (CD45.1) mice 4 days after immunization with soluble TM4 core or N276D Env. Numbers in the plots indicate % of cells in the CTV high gate. **(B)** Quantitation of data in A. Data are representative of two independent experiments analyzed days 3-4 post-immunization (n=3-4 mice per group). ****, P < 0.0001. Ordinary one-way ANOVA (Tukey).

**Supplementary Table 1. B cell transfer efficiency.** Transfer efficiency was determined from 3 mice per dilution group as described in Materials and Methods.

**Supplementary Table 2. Antigen nomenclature.** Properties of soluble envelope trimer antigens. Affinities were determined using the BLI assay as described in Materials and Methods.

